# Pharmacophoric-based ML model to filter candidate E3 ligands and predict E3 Ligase binding probabilities

**DOI:** 10.1101/2023.08.10.552794

**Authors:** Reagon Karki, Yojana Gadiya, Simran Shetty, Phillip Gribbon, Andrea Zaliani

**Affiliations:** Fraunhofer Institute for Translational Medicine and Pharmacology (ITMP), Schnackenburgallee 114, 22525 Hamburg, Germany; Fraunhofer Cluster of Excellence for Immune-Mediated Diseases (CIMD), Theodor Stern Kai 7, 60590 Frankfurt, Germany; Bonn-Aachen International Center for Information Technology (B-IT), University of Bonn, 53113 Bonn, Germany; Hamburg University of Applied Sciences (HAW), 20099 Hamburg, Germany

**Keywords:** E3 Ligase ligands, Machine Learning, Virtual Screening, Protein degradation, ErG fingerprint, PROTACS, Compound Libraries

## Abstract

Among the plethora of E3 Ligases, only a few have been utilized for the novel PROTAC technology. However, extensive knowledge of the preparation of E3 ligands and their utilization for PROTACs is already present in several databases. Here we provide, together with an analysis of functionalized E3 ligands, a comprehensive list of trained ML models to predict the probability to be an E3 ligase binder. We compared the different algorithms based on the different description schemes used and identified that the pharmacophoric-based ML approach was the best. Due to the peculiar pharmacophores present in E3 ligase binders and the presence of an explainable model, we were able to show the capability of our ErG model to filter compound libraries for fast virtual screening or focused library design. A particular focus was also given to target E3 ligase prediction and to find a subset of candidate E3 ligase binders within known public and commercial compound collections.

## Introduction

E3 Ligases are a group of hundreds of enzymes that play a crucial role in ubiquitination, a post-translational modification that regulates protein turnover, signal transduction, DNA repair, and many other cellular processes^1^. By ubiquitination, the E3 Ligase catalyzes the transfer of ubiquitin, a small tagging protein, to a substrate protein, marking it for degradation by the proteasome or altering its activity, localization, or interactions with other proteins. Based on this ubiquitination mechanism, E3 Ligases are classified into two prominent families: the RING (Really Interesting New Gene)^2^ and the HECT (Homologous to E6AP C-Terminus)^3^. The former, conserved across species such as yeasts, demonstrates an internal ubiquitin mechanism upon activating E2-enzyme Cdc3 by forming a complex with Cub-Rbx1/Roc1 (sub-domains of the RING family)^4^ while the latter ubiquitinates via the classic thioester mechanism^5^. Furthermore, individual E3 Ligases display specificity towards distinct substrates through their diverse domains or subunits, enabling them to interact and function effectively in various cellular environments^6^. Their crucial regulatory role revolves around ensuring a balanced protein composition within cells and safeguarding against the buildup of misfolded, damaged, or abnormal proteins that could lead to diseases. Indeed, mutations or dysregulation of E3 Ligases have been associated with various disorders, including cancer, neurodegeneration, inflammation, and viral infections^7–9^. As a result of this, E3 Ligases have emerged as attractive therapeutic targets, especially in cancer, leading to several clinical candidates as inhibitors of E3 Ligases^10^.

The diversity and complexity of E3 Ligases affect the development of selective and potent inhibitors of E3 Ligases. This necessitates a comprehensive understanding of the particular E3 Ligase in question, including its substrate specificity, binding site traits, and structural attributes. Nevertheless, the discovery of potential ligands for E3 binders holds significant importance for several reasons. Firstly, it can assist in deciphering the regulation mechanism employed by E3 in different contexts (i.e., in regulatory or diseased conditions). Identifying ligands that bind to specific domains or interfaces of E3 Ligases can reveal their structural and functional features, helping elucidate molecular mechanisms of substrate recognition, ubiquitin transfer, and regulation^1,11,12^. This knowledge can facilitate the design of more selective and effective inhibitors of E3 Ligases. Furthermore, exploring the interaction mechanisms between E3 Ligases and other proteins or small molecules can lead to the identification of novel substrates and shed light on their roles in cellular signaling and processes^13^. Secondly, discovering ligands capable of regulating the function or translocating E3 Ligases, thereby manipulating their substrate specificity and selectivity, can serve as novel therapeutic targets and strategies across numerous indication areas^14,15^. For example, ligands that induce the degradation of disease-causing proteins or activate the immune response are explored as potential treatments against cancer^16,17^. Thirdly, this discovery marks a shift in traditional drug development, moving towards a new era where intracellular mechanisms are activated rather than relying solely on external agents. In such cases, the ligands that bind to E3 Ligases can serve as starting points for drug discovery campaigns, thus helping in the optimization of the properties and efficacy of lead compounds^18^. Furthermore, ligands that compete with or allosterically modulate the binding of other proteins or small molecules to E3 Ligases can provide insights into their functional and regulatory networks^19^. Discovering new ligands for E3 Ligases is crucial to understanding their complex roles in cellular homeostasis and developing effective therapies that harness their potential for targeted protein degradation and immune activation.

So far successful attempts to identify novel E3 Ligase ligands have been through 1) synthetic chemistry: libraries of compounds can be assembled and screened for binding affinity to the E3 Ligase of interest^20^; 2) computational methods such as molecular docking and virtual screening^21^; and 3) fragment-based structural biology approaches^22^. In either of the above-mentioned cases, a compound library is a prerequisite following in-vivo or in-silico validation of potential E3 Ligase ligands. We leverage the results from these techniques to formulate a collection of proven E3 Ligase binders and attempt to rationalize it^23^. In this manuscript, we constitute an overview of the chemical space spanned so far by E3 Ligase binders, comparing it with the chemical space of known non-binders. By doing so, we define filtering possibilities to efficiently scan commercial ligand libraries to be eventually tested and generate hypotheses on the previously unproven mechanism of actions for old approved or withdrawn drugs, like thalidomide analogs, which can be eventually fruitfully used as E3 Ligase binders. Additionally, we present a Machine learning (ML) model that anticipates the E3 Ligase classification by utilizing the Extended Reduced Graph (ErG) approach, a 2D pharmacophoric fingerprint. Such a model holds significant potential for refining virtual compound libraries or segmenting commercial compound collections with a specific focus.

## Methods

### Dataset Construction

The unique compounds with proven E3 binder nature have been either taken from PROTAC-DB 2.0^24^ and PROTACpedia^25^ collections or generated from Protein Degrader Database (PDD)^26^ **(Table 1)**. To enhance the numerosity of this set, we added published E3 binders (not registered as PROTACs), like the DCAF1 binders which have been published with extensive SAR^27^. With this, we have compiled the largest E3 Ligase binder collection to date. For negative samples i.e. non-E3 ligand binders, we selected compounds publicly available in ChEMBL (version 33) which reported binding affinities equal to or above 100uM (with pChembl ≤ 4) under any bioassay type (i.e., binding, functional, ADME, toxicity) against human targets^28^. To assure the quality of data used, we selected assay reports that showed the highest manual annotation score at confidence level 9. This strategy should prevent our selection from collecting molecules with high affinity E3 binders as so far reported. Moreover, the “non-E3 ligand binders” chemical space so sampled should be more diverse and cover different pharmacophoric patterns. Of course, these criteria have to be considered as dynamically changing as this dataset grows daily with reported scientific and patent literature and needs to be constantly updated. A summary of the collection is provided in **Table 1**. Finally, the descriptors generated by these molecular structures have been made public to ensure the reuse and reproducibility of the data and models respectively.

**Table 1:**
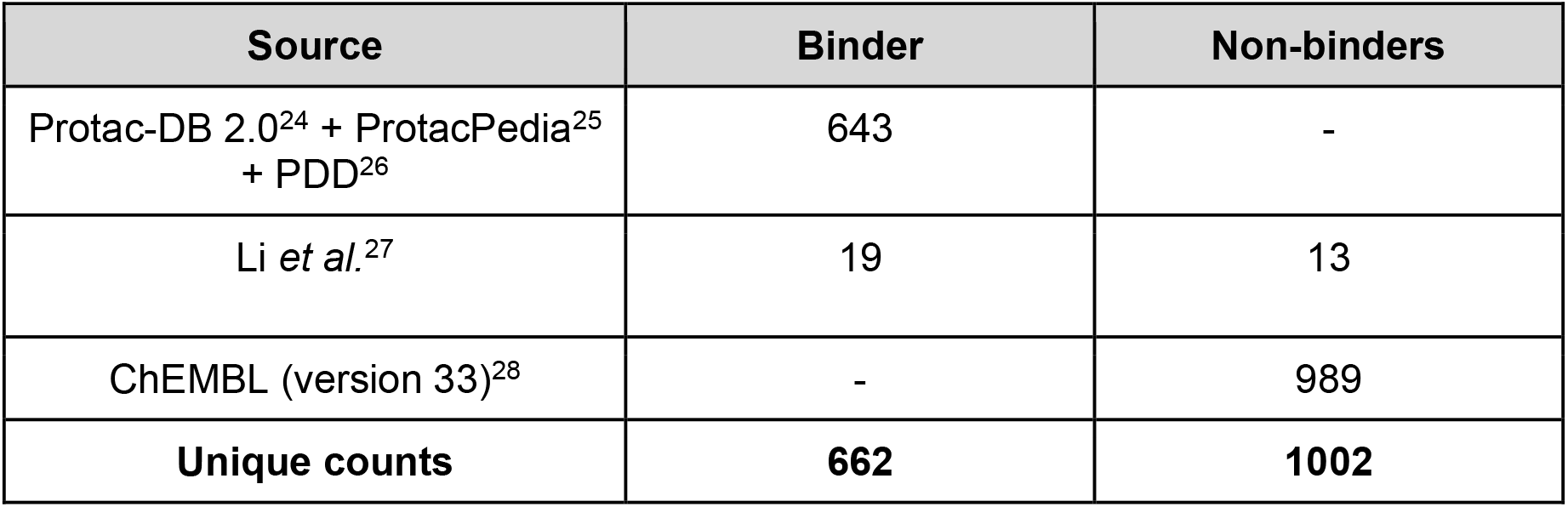
Overview of the E3 Ligase binders/non-binders dataset.

| Source | Binder | Non-binders |
| --- | --- | --- |
| Protac-DB 2.0<sup>24</sup> + ProtacPedia<sup>25</sup><br>+ PDD<sup>26</sup> | 643 | - |
| Li *et al.*<sup>27</sup> | 19 | 13 |
| ChEMBL (version 33)<sup>28</sup> | - | 989 |
| **Unique counts** | **662** | **1002** |

### Molecular Description

The collected molecule structures, after the removal of salts, neutralization, and removal of duplicated InChiKeys, have been used to generate thirteen classical chemico-physical descriptors, like LogP, molecular weight (MW), polar surface area (PSA), etc. We refer to these descriptors as ChemPhys in the remaining manuscript. We also calculated for each molecule feature vectors in the form of circular Morgan^29^ (ECFP4; radius = 4), topological RDKit^30^, smart-based MACCS^31^, and path-based Avalon^32^ fingerprints through RDKit Library and the ErG pharmacophoric fingerprint^33^ as implemented within MOE 2022.02 (CCG group)^34^ through the relative publicly available script^35^. While the classic fingerprints like MACCS, Morgan, Avalon, and RDKit are known and commonly used within the chemoinformatic community, ErG fingerprints have received less attention in recent years^36^. They are count-based 315 bits long fingerprint vectors and provide a very compact description of molecules. The ErG scheme associates the molecule 2D structure with six different classical pharmacophoric entities (i.e. H-bond donor, H-bond acceptor, hydrophobic group, aromatic ring system, and positive/negative charge) and counts through their shortest path how many reciprocal bond distances are present. So, for instance, an ErG bit called Ar_Hf_d4 will count how many times in a molecule the centroid of an aryl moiety (Ar) distant 4 bonds from a hydrophobic moiety (Hf) is encountered. We were aware that this scheme is not exempt from limitations or redundancies such as the standing out of Ac_D_3 descriptor for the repeating amido groups in peptides or no discrimination between the meta and para phenyl substituent positions as both have the same distance of at least 2 from any other pharmacophoric point attached to phenyl at position 1. Nevertheless, we decided to use this unusual description among other fingerprinting schemes, as the ErG scheme allows easy clusterization of different molecules providing the same pharmacophore structure, fast calculations, and explainable meaning of each bit, as shown in **Figure 1**.

**Figure 1:**
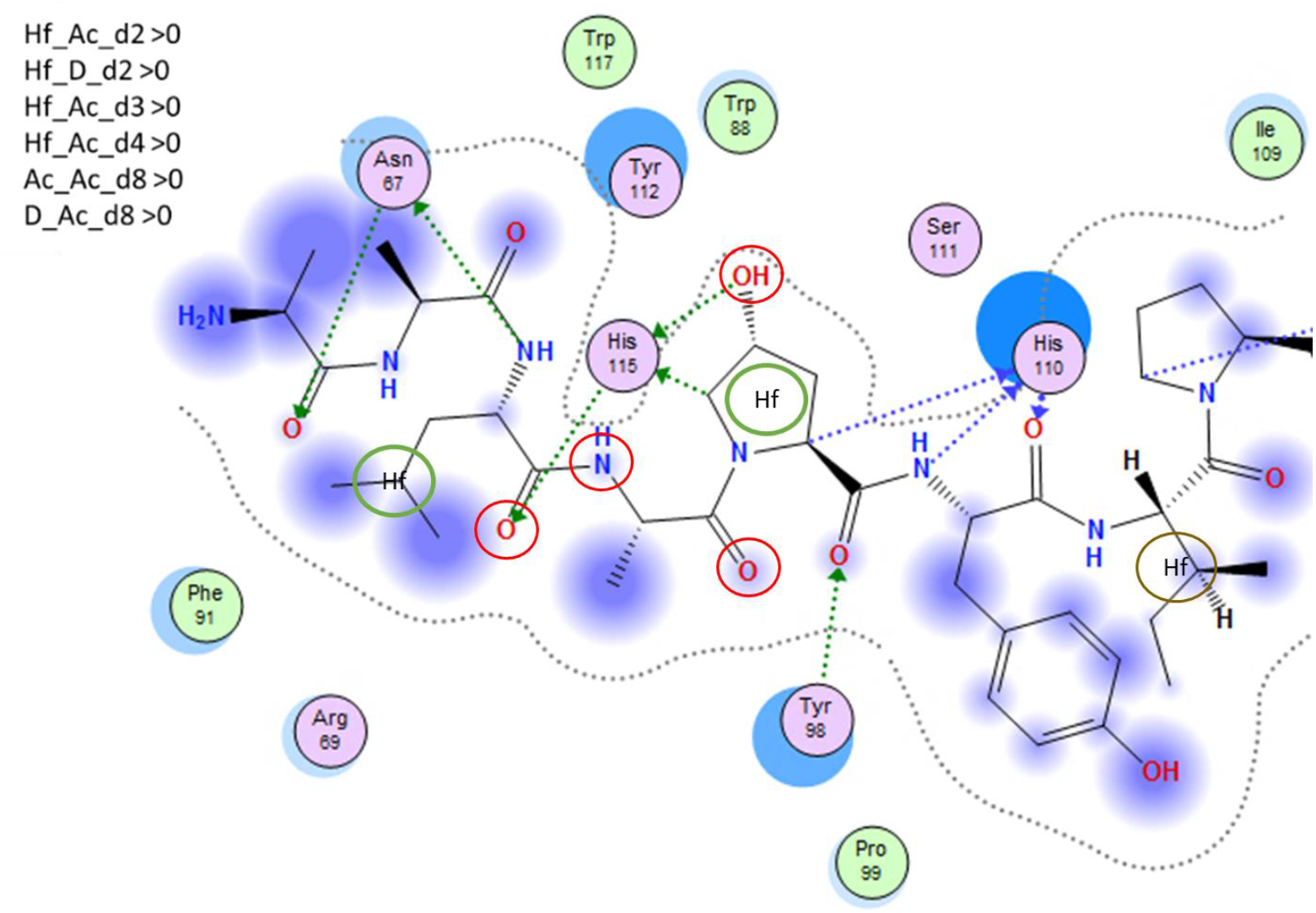
Interpreting pharmacophoric interactions from ErG fingerprints. The ligand-protein interactions from PDB for von Hippel-Lindau - Elongin B - Elongin C complex bound to HIF1-alpha peptide (PDB:4AJY) show non-null ErG bits that can be easily extracted from six pharmacophoric atoms: three acceptors N, O, OH [Ac] (circled in red), two donors O, OH [D] (circled in red), and two hydrophobic centers [Hf] (circled in green).

We optimized the ErG fingerprints by restricting the maximum distance to 15, thereby limiting the length of the ErG vector to 315 (21 pairs X 15 distances). A fuzzy component (of 0.3) was also added to the ErG bit count, meaning that each ErG bit count localized at side distance (i.e. distance +/-1 bond), was augmented by 0.3 units. In this context, a distance is counted through the shortest path between the bonds in a 2D graph. Thus, when we filtered the dataset using Hf_Ac_d2 > 0.3 or Hf_D_d2 > 0.3, we ended up with 463 compounds, of which 446 are known E3 Ligase binders against only 17 non-binders **(Figure 2)**. This demonstrated the usefulness of an intuitive fingerprint scheme like ErG as we were able to enrich the E3 binders selection up to 96% with just two of the six bits suggested for VHL binders.

**Figure 2:**
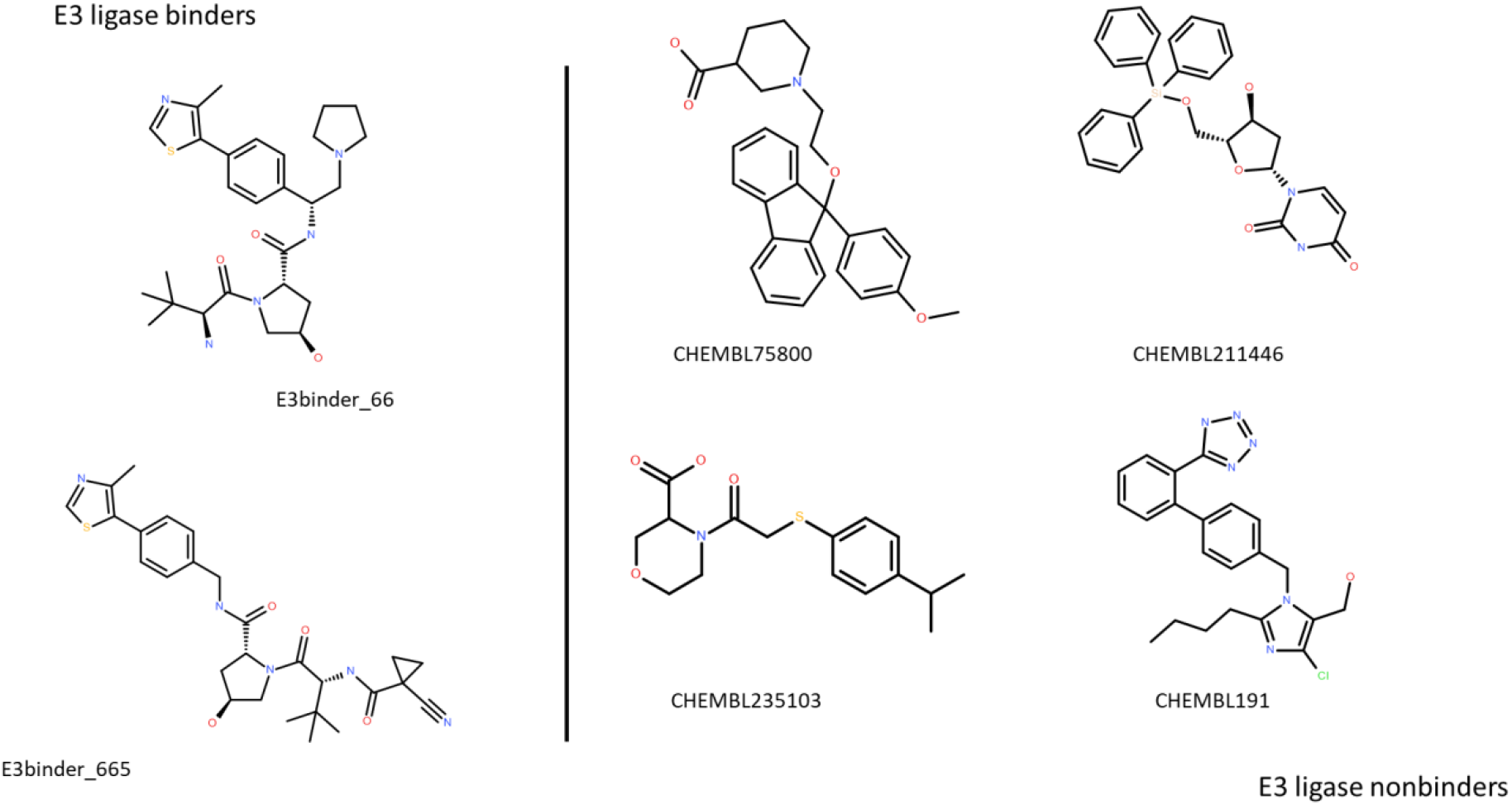
Example of molecules retrieved from the dataset using the two ErG bit selections (Hf_Ac_d2 > 0.3 or Hf_D_d2 > 0.3)

### Structural distinctions within the dataset

In order to evaluate structural overlap within the E3 binder dataset, we generated a fragmentation tree using the MHFP6 fingerprints^37^ in a TMAP based visualization (https://github.com/reymond-group/tmap) **(Figure 3)**. MHFP6 is a circular fingerprint, similar to ECFP4, which utilized the nearest neighbor-based mini hashing thus allowing in identifying analogs of compounds^38^. Each branch in the self-organizing TMAP is generated by analogs with the same common scaffold. From the map, no singletons or isolated branches were found. Compounds are linked by the presence of mutually similar substructures. Furthermore, it is interesting to note that orange dots (E3 Ligase binders families of analogs) are sparsely distributed within the tree, even if the majority are aggregated in the bottom left part of the plot.

**Figure 3:**
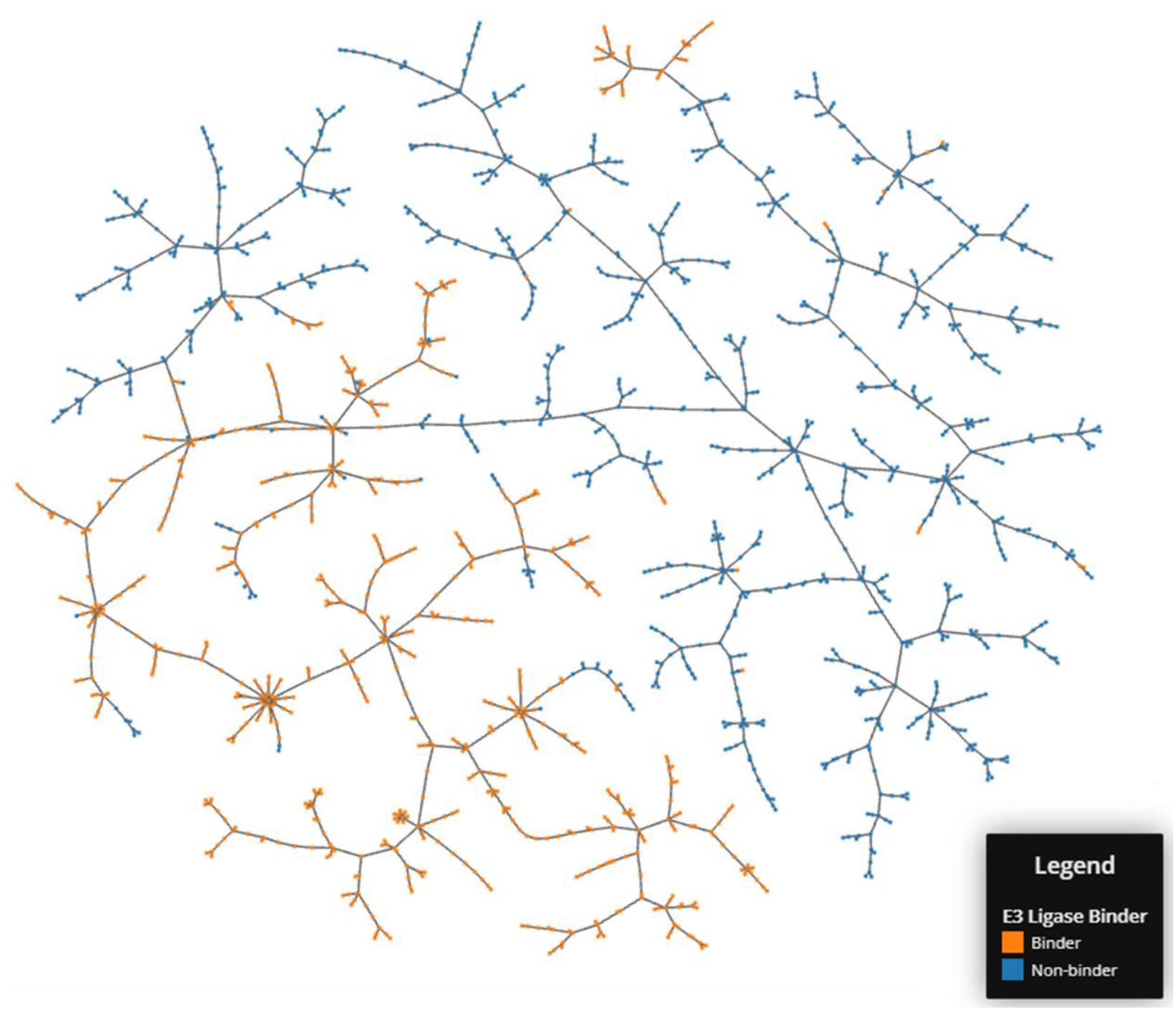
The fragment TMAP tree connecting structural families of the dataset. Orange dots correspond to known E3 binders, while blue ones denote molecules without E3 Ligase binding capabilities.

### Machine learning models

The KNIME 4.7.2 platform was used to build the ML classification models and building the dataset for the models. The KNIME ML engine^39^ (AutoML node) generated several models with different classification algorithms namely logistic regression (LR), naive bayes (NB), random forest (RF), support vector machines (SVM) and XGBoost. The KNIME node automatically selects the best algorithm on the basis of the highest Cohen’s kappa^40^ value defined in here below:

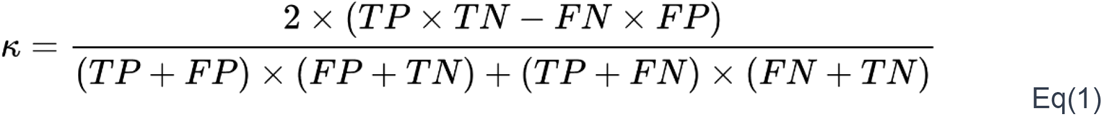

Where TP and TN refer to true positive and true negative predictions respectively, while FP and FN refer to false positive and false negative predictions respectively.

The descriptors matrix used in each model has been reduced removing columns possessing variance higher than 0.2. In doing so, no columns were removed for the ChemPhys matrix, while the ErG description matrix was reduced to almost half. For the selection of the best model, a 4x cross-validated strategy was implemented. Additionally, the influence of each descriptor on the model was evaluated using KNIME’s “Global Feature Importance” component^41^. Upon selection of the best model, this model is optimized through a random search involving different parameters such as learning speed (eta range between 0.2 and 1) and number of epochs (range between 1 and 10) with inner cross-validation (10x) strategy.

We also report the performance of the model using two commonly known metrics, accuracy and Area Under the Receiver Operating Characteristic curve (AUROC). Accuracy is commonly used for balanced datasets to understand how good a model is at predicting the true labels (true positives and true negatives), and the AUROC is used for imbalanced datasets to understand how well the model can discriminate one class from another.

## Results

### Identification of structural clusters within E3 binders and non-binders

Before starting to model the classification labels E3 Ligase “binder/non-binder”, we explored the dataset we collected by means of “classical” statistical algorithms to check whether certain groups or particular clusters emerged. To do so, we performed PCA analysis **(Supplementary Figure 1S)** and represented the ErG descriptor t-SNE **(Figure 4)**. The PCA analysis highlighted how Cereblon (CRBN) and Von Hippel-Lindau (VHL) binders can be easily clusterized through the ErG pharmacophoric fingerprint scheme. Analogously, in the t-SNE plot, these two major groups of E3 Ligases are clearly distinct. We also found a small cluster in the t-SNE plot that corresponded to CRBN binder analogs containing N-alkylated succinimide moiety. Their cluster separates nicely from the CRBN mother group possibly due to their considerably larger size than non-N-alkylated succinimide-containing CRBN binders. Also, indisulam-like sulphonyl-containing E3 binders were found to be in a separate cluster.

**Figure 4:**
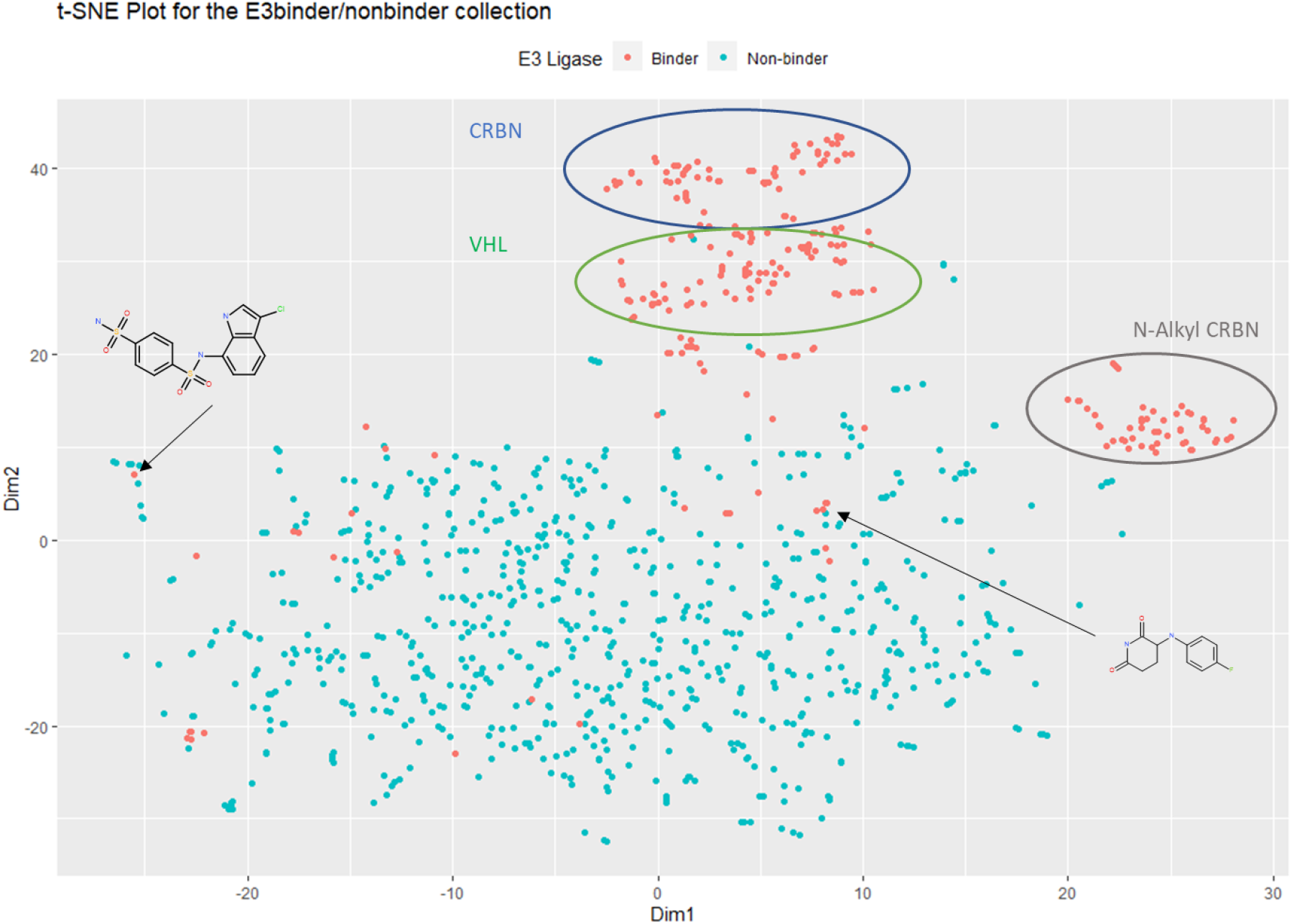
t-SNE plot of the Erg descriptor matrix. Certain groups are clearly recognizable according to relative E3 Ligase targets (CRBN and VHL).

### Explainable model assists in equating ErG performance with other fingerprints

Several general chemoinformatic analyses have been recently published from Protac-DB molecules only and their building blocks^42,43^, however, as far as we know, none trying to map the pharmacophoric content of this chemical space. Our statistical machine learning models successfully predict the probability of being an E3 Ligase binder with high accuracy **(Table 2)** and significant AUC-ROC **(Supplementary Figure 2S)**. This led to two key outcomes: first the Extended Reduced Graph-based (ErG) fingerprinting performed comparable to the other fingerprints and second, this ease of interpretability of the pharmacophoric models made them a suitable choice for future explainable artificially intelligent (XAI) algorithms. While there might be a contention that alternative fingerprints, distinct from ErG, rely on a certain category of molecular substructure pattern that could be linked to particular pharmacophoric attributes, they remain less intuitive compared to the ErG-derived fingerprinting approach. ErG fingerprint was found to outperform the classic circular ECFP4 fingerprints indicating that a specific pharmacophoric description is an optimal way to describe E3 binder molecules in a way dependent on distinctly located interacting atoms rather than on specific substructures. In other words, ErG description was found to be more inclusive than ECFP4 in terms of atom sets needed for interactions. Only a higher number of “useful” bits rescued the modeling performances of more structural fingerprints like Avalon and RDKit.

**Table 2:** Performance summary of the fingerprints and chemphys description matrix. The result was reported on the external test set (20% of the total dataset). The number of bits reported in the table is the final bit count post-filtering as mentioned in the Methods section.

| Fingerprint | Best Model | N.Bits used | Accuracy | Cohen’s Kappa | AUC-ROC |
| --- | --- | --- | --- | --- | --- |
| ChemPhys | XGBoost | 13 | 0.973 | 0.943 | 0.995 |
| Avalon | XGBoost | 257 | 0.976 | 0.95 | 0.994 |
| RDKit | XGBoost | 901 | 0.976 | 0.95 | 0.993 |
| ECFP4 | XGBoost | 30 | 0.937 | 0.866 | 0.974 |
| **ErG** | **XGBoost** | **83** | **0.964** | **0.924** | **0.980** |

Additionally, we compared the models using the ChemPhys feature to compare it against the model using the ErG descriptor. We found that both models show an equal model specificity (0.98) while looking at the confusion matrix of ErG **(Supplementary Figure 4S)** with ChemPhys **(Supplementary Figure 4S)**. We also looked into the distribution of the 13 ChemPhys properties in separating the binders from non-binders **(Supplementary Figure 6S)**. All the features that were used showed significantly different between the binders and non-binders.

### Pharmacophoric features reveal characteristics of E3 ligase binder

Next, we looked into the range of explainable features from the compound structure perspective (found in chemo-physical) that could provide hints for distinguishing the E3 binders from non-binders. We found that the presence of amide bonds in the molecule is an important feature in identifying E3 binders **(Supplementary Figure 3S)**. Although the amido groups or indisulam-required sulfonyl groups contributed to raising the polar surface of the compound, it hampered the chance of central nervous system (CNS) penetration, a known problem for several PROTAC compounds^44^. These findings, however, illuminate that CRBN binders as succinimides play a fundamental structural motif in binding. Moreover, the presence of amide groups, while necessary in CRBN binders, might not be of importance when dealing with peptidomimetic VHL binders. Hence, we should not take these chemo-physical descriptions as “rules” indicative of an E3 ligase binding as they could be specific only in certain cases. The ErG-derived pharmacophoric “rules”, instead, were found to be more robust, useful, and detailed across different ligase binders.

When looking into the top ten most influential ErG bits reported by the XGBoost model, it appears that Hf_Ac_d2 is on the top followed by Hf_Ar_d6 **(Figure 5)**. These two bits capture pairs of hydrophobic (Hf) patches of at least three carbon atoms and acceptors (Ac) or aromatic centers (Ar) at a distance of 2 and 6 respectively. Our finding revealed that the top-performing bits, Hf_Ac_d2 and Ac_Ac_d6, are present in the succinimide ring, while the hydroxyproline moiety, which is a crucial residue for VHL binders, only has Hf_Ac_d2. This level of local interpretability has no comparison with any other fingerprint methods in use; and to have the same level of explanation we should use “back-projection” of singular bits onto chemical structures to explain, for instance, the relevancy of a precise and relevant circular fingerprint bit^45^. Thus, we demonstrated that the ErG bits are driving recognition of E3 ligase binders by highlighting the most relevant of them for known ligase-ligand interactions.

**Figure 5:**
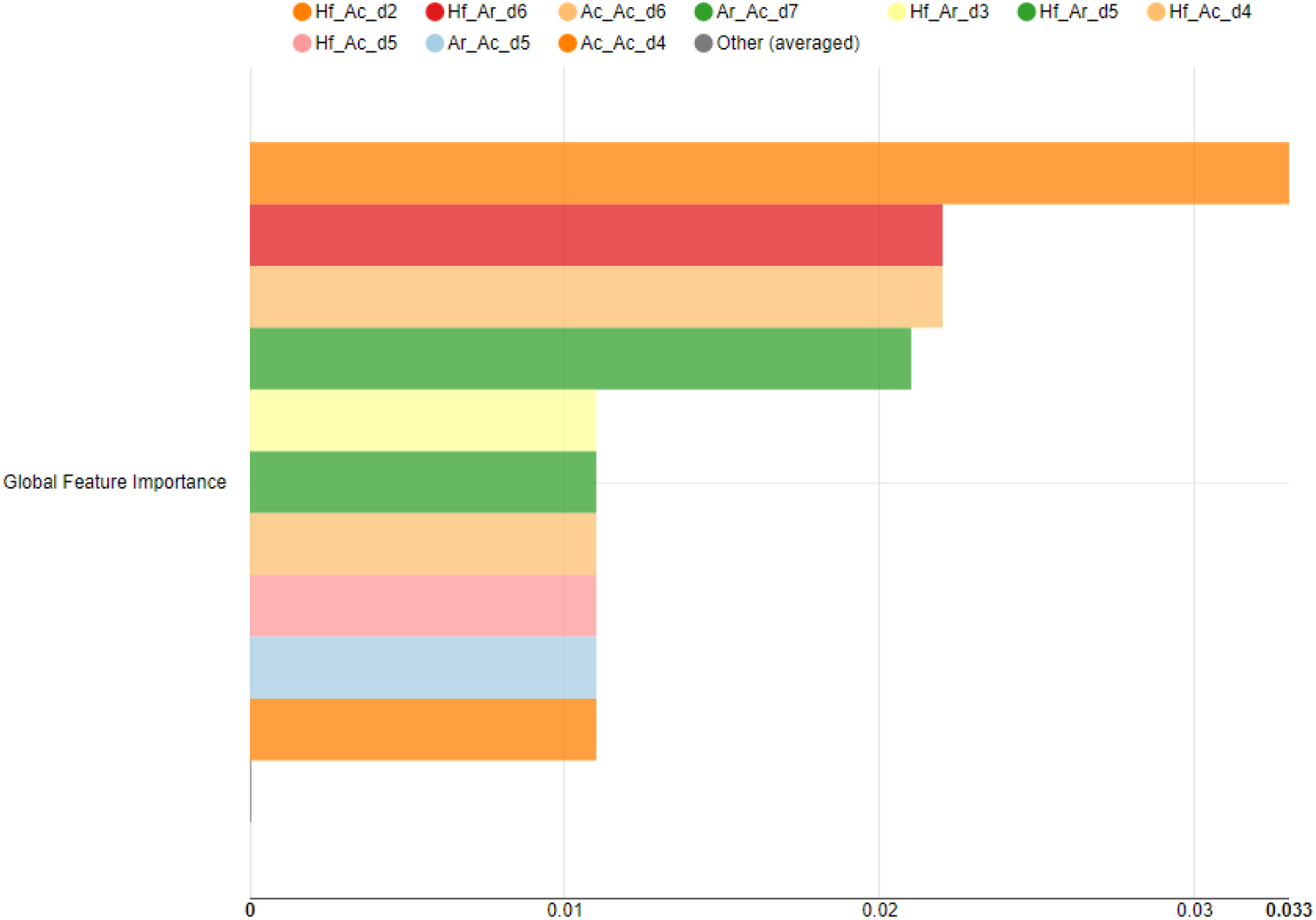
Variable importance in XGBoost model with ErG description. Feature importance is calculated by counting how many times a feature (ErG Bit) has been selected for a split and at which rank (level) among all available features (candidates) in the trees of the random forest. A higher value indicates higher feature importance. The ErG bit names were coded as follows: e.g. Hf_Ac_d2 means Hydrophobe_Acceptor_at_bond_distance_2 in the molecule’s reduced graph.

A more precise analysis is depicted in **Figure 6**, where the top nine “relevant-to-model” ErG bits have been profiles through boxplots and their reciprocal mean difference significances. Through this, we can extend the usage of these filtering rules to model E3 ligase families and, possibly, their pharmacophoric requirements. Recently, Karki *et al*. (2023, accepted) have published their work in this direction trying to get a model to predict the most probable E3 ligase for the binders^46^.

**Figure 6:**
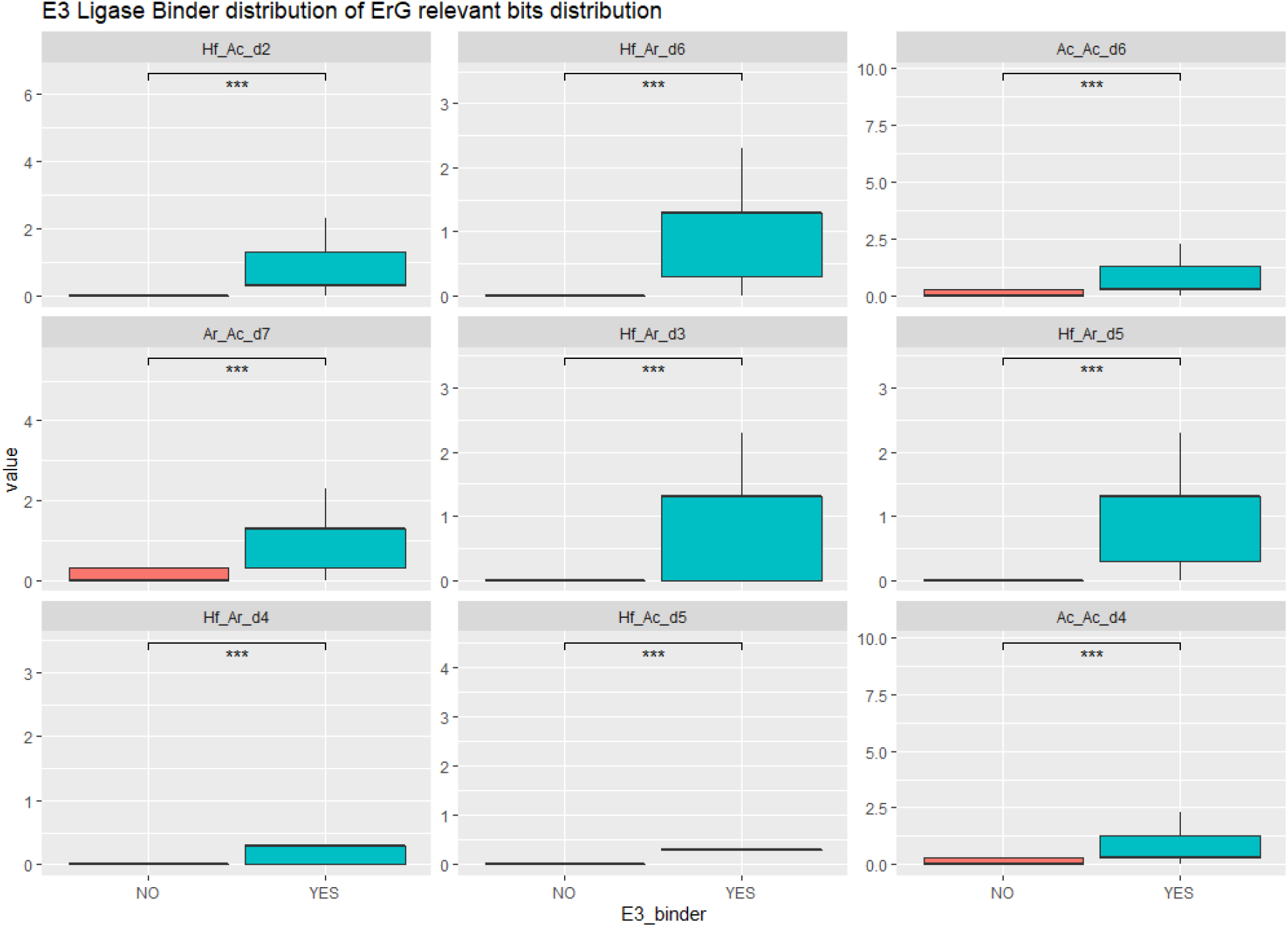
Variable Distribution of most relevant ErG bits. Boxplots of the top nine most relevant ErG bits. All ErG bits shown are statistically significant with respect to their mean differences (*** = p-value < 0.05).

### Discovery of novel E3 ligases from compound libraries

To validate the predictions of the model, we thought to predict external molecular sets, but with published evidence of E3 ligase binding properties. We chose a newly published set of E3 ligase zinc finger protein, atypical E3 Ubiquitin Ligase (ZFP91) binders^47^, where only Napabucasin among all the molecules described in supplementary material was not predicted as E3 Ligase binder **(Supplementary Table S1)**. Next, we assessed the compound’s similarity to the training dataset. Out of the 32 molecules sampled, eight compounds, apart from Napabucasin, demonstrated a Tanimoto distance above 0.5. All these compounds were predicted as E3 binders by our model. Since ZFP91 binders are novel and were not represented in our dataset, at present, Napabucasin represents an outlier for our model.

Another application case we present here is to predict E3 binders from small compound libraries either publicly or commercially available. We took the commercial Asinex libraries^48^ dealing with CRBN binding or with molecular glues as targets and merged them into one dataset. Out of the resultant collection of 1257 compounds, 199 (16%) were predicted as E3 binders. This prediction, however, drops if similarity to training above 0.5 Tanimoto is taken into account: 42 (22%) compounds predicted as E3 binders out of 193 remaining (15%). We also tested the Fraunhofer ITMP Repurposing Library (ca 8.5K compounds)^49^, where out of 2500 compounds having Tanimoto similarity to training higher than 0.5, 85 compounds were predicted as E3 binders (0.9%). Of these predicted E3 Ligase binders Glutethimide, a known sedative, with a binding of 39uM ^50^ to CRBN, and Idasanutlin, a MDM2 Ligase binder^51^, were easily identified and deserve more attention. We have also predicted known repurposing collections, like ENZO^52^ and LOPAC^53^ **(Table 3)**.

**Table 3:** Summary table for number of E3 ligases predicted from known compound libraries.

| Collection name (version) | No. of compounds | No. of predicted E3 Ligase binders | Predicted E3 Ligase binders (%) |
| --- | --- | --- | --- |
| Broad (2017) | 5632 | 320 | 5.7 |
| Enzo (v.3) | 764 | 62 | 8.1 |
| Lopac | 1240 | 31 | 2.5 |
| Fraunhofer ITMP Repurposing | 8494 | 549 | 6.4 |

## Discussions

Filtering commercial collections of compounds through machine learning predictions offers several advantages. In terms of efficiency, ML algorithms can rapidly screen large databases of compounds, filtering out those that are unlikely to be useful for a particular application. In terms of practical cost savings, filtering out compounds that are unlikely to be useful can reduce the cost of purchasing and testing large numbers of compounds. This is particularly important for drug discovery, where the cost of synthesizing and testing compounds can be high^54^. In terms of enrichment of active compounds (hit rate increase), researchers can improve their chances of identifying compounds that are likely to be effective for a particular application. At the moment very few chemical vendors, as far as we know, compiled commercial E3 ligase library collections, and if they did, collections are limited to analogs of known CRBN or VHL binders. In terms of reduced experimental variability, ML algorithms can help to identify compounds with similar properties, which can reduce experimental variability and increase the possibility to surface real quantitative structure-activity relationship (QSAR) which can then be used to refine prediction models and support more robust prioritization of scaffolds. Finally, in terms of structural novelty, pharmacophoric filtering might help to identify compounds that allow access to novel chemical space and generate IP.

Using the largest dataset of experimentally validated E3 ligase-substrate interactions, the model presented here was trained to identify key features that influence the binding probability and selectivity of E3 Ligases. We generated several models all with excellent performances and only limitations of the number of molecules in both classes (E3 binders and non-binders) might limit the generalizability of the pharmacophoric information found in this analysis. However, the fact that the ErG description can clearly generate rules for the E3 ligase specificity is an encouraging finding which opens up several filtering future opportunities for a fast and efficient virtual screening of enormous commercial molecule collections.

### Does a common recognition motif exist for E3 ligases?

Some E3 Ligases share common features in their ligand-binding cavities that can be exploited for drug discovery. For example, many RING-type E3 ligases have a conserved cysteine residue in their catalytic site, forming a covalent bond with ubiquitin during ubiquitin transfer^55^. This cysteine residue can be targeted by electrophilic or covalent inhibitors that mimic the structure of ubiquitin and bind to the active site of the E3 ligase (PDB:5EDV). Similarly, some E3 ligases have specific domains or motifs that mediate protein-protein interactions and substrate recognition. For example, the VHL tumor suppressor protein, a HECT-type E3 Ligase, contains a hydrophobic pocket that binds to the hydroxyproline residue of the hypoxia-inducible factor (HIF) transcription factor, marking it for degradation under normoxic conditions (PDB:1LM8)^56^. This pocket has been targeted by small molecule inhibitors that prevent HIF binding and stabilize HIF, offering a potential therapeutic strategy for cancer. Other E3 Ligases, such as the Cullin-RING Ligases (CRLs), have more complex structures and interactions with substrate adaptor proteins making their ligand-binding cavities challenging to target^57^. This field of study is growing day by day^21,43,58,59^. Finally, CRBN, since 2018, has inspired researchers to find the proper chemical to bind this important E3 Ligase^50^.

We are aware of the dynamic nature of the field: each novel ligase ligand should be added to the training set to improve the generality of the model. Moreover, inactive non-binders structurally related to the active ones should be added as well in order to reduce the diversity gap, which might drive the actual models. Nevertheless, we took several steps to address this potential issue. First, we curated the dataset to include only high-quality literature data. Second, we performed rigorous cross-validation to evaluate the generalizability of the model to unseen data. Third, we used feature selection techniques to use the most informative features that contribute to binding probability to E3 ligases.

Finally, we validated the model on an independent test set and observed excellent performances, indicating that our models were not simply memorizing the training data. We think that our results provide valuable insights into the compounds that demonstrate E3 ligase binding capabilities.

## Data availability

All input data and workflow have been made available on GitHub at https://github.com/Fraunhofer-ITMP/E3_binder_model_paper2.

## Author contribution

AZ and RK conceived the work. RK programmed mapping of the E3 binders in accordance with the three database sources cited. AZ, YG, and SS performed the analysis and contributed to ideation. AZ, YG, PG, and RK have written the manuscript. All the authors have read and approved the final manuscript.

## Acknowledgments

We acknowledge PROTAC-DB and PROTACpedia curators for the access to PROTACs data and Dr. Aniket Ausekar from Evolvus inc. for providing license to access PDD data.

