## Supplementary Files for "Pharmacophoric-based ML model to filter candidate E3 ligands and predict E3 Ligase binding probabilities"

### Contents

- **Supplementary Figure 1S:** Two component PCA plot from ErG dataset
- **Supplementary Figure 2S:** AUC-ROC curve for XGBoost model with ErG description
- **Supplementary Figure 3S:** Distribution of most relevant ChemPhys features
- **Supplementary Figure 4S:** Confusion matrix of the test set using ErG descriptor
- **Supplementary Figure 5S:** Confusion matrix of the test set using ChemPhys
- **Supplementary Figure 6S:** Boxplots for ChemPhys description
- **Supplementary Table S1:** Table for compounds with SMILES predicted as E3 ligase binders

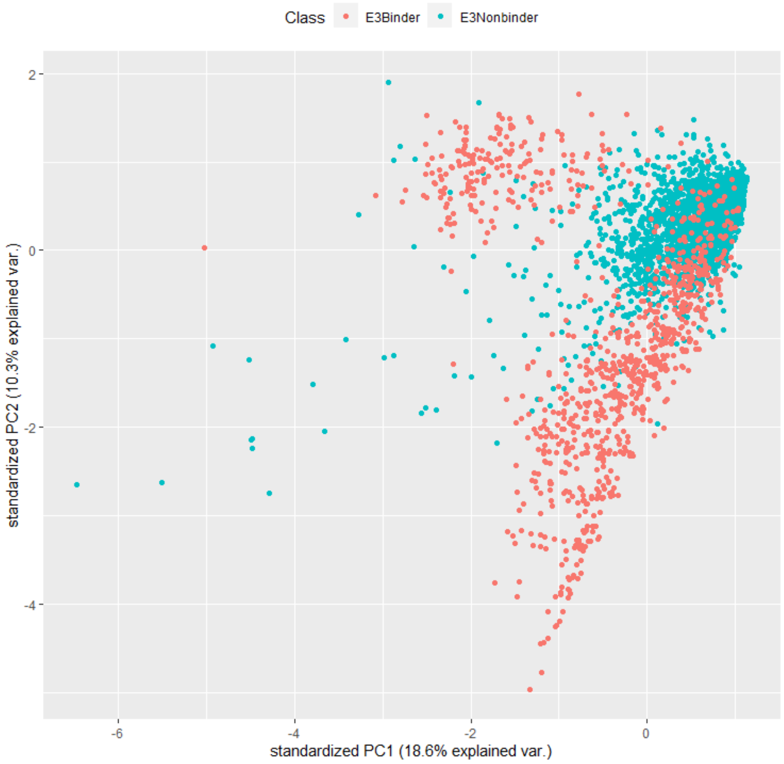


**Supplementary Figure 1S:** Two component PCA plot from ErG dataset. The figure shows two well-separated clusters one in red highlighting major target VHL and CRBN and other in blue highlighting non-binders.

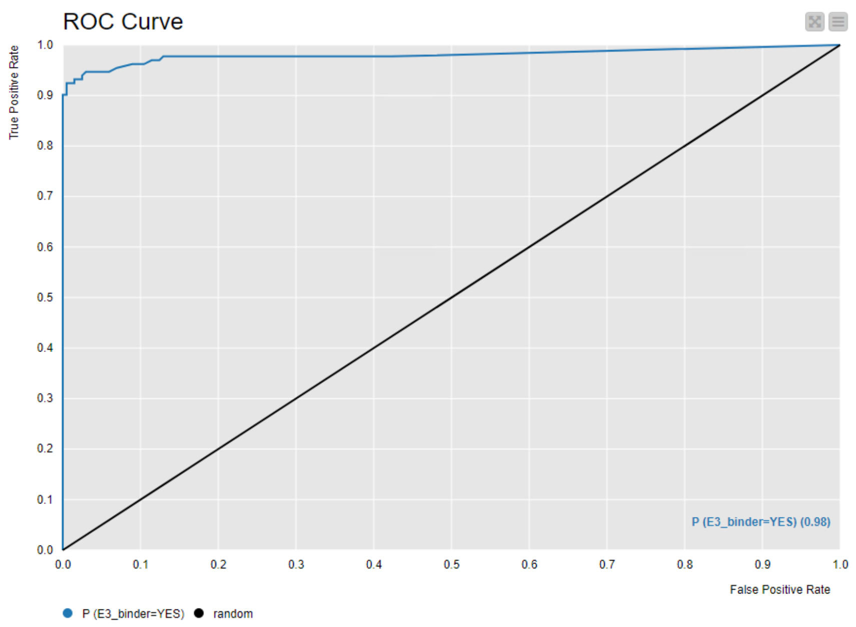


**Supplementary Figure 2S:** AUC-ROC curve for XGBoost model with ErG description.

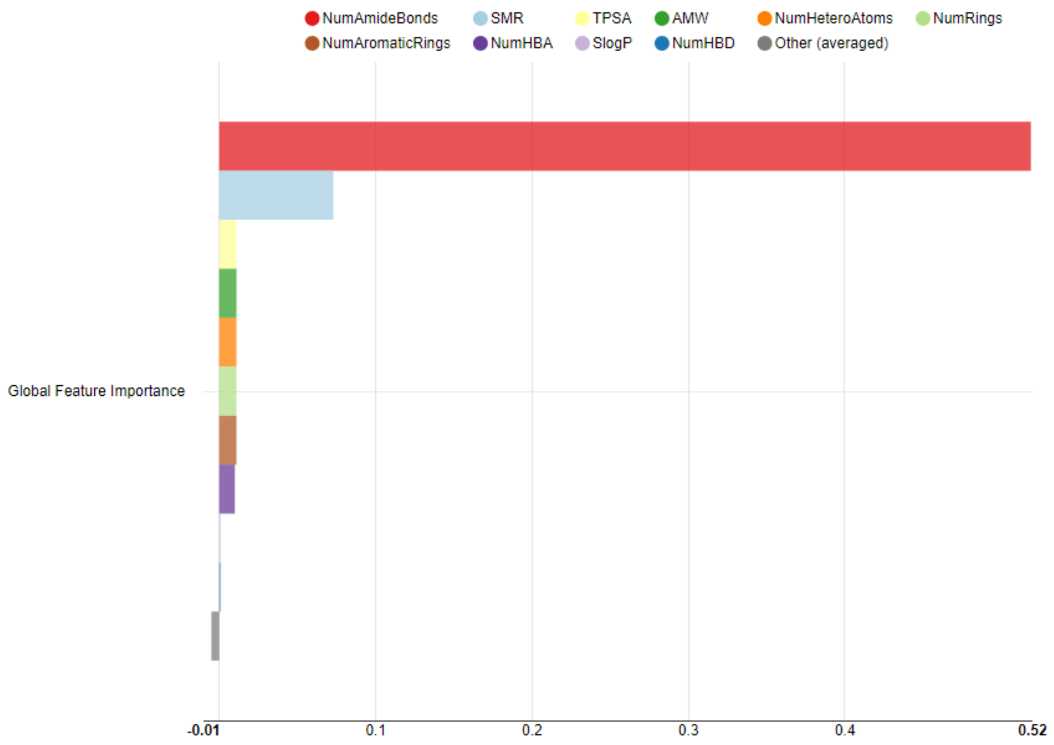

**Supplementary Figure 3S:** Distribution of most relevant ChemPhys features. The number of amide bonds is found to be the differentiator of the distributions with highest importance and this is a basic structural bit component in both VHL and CRBN binders.

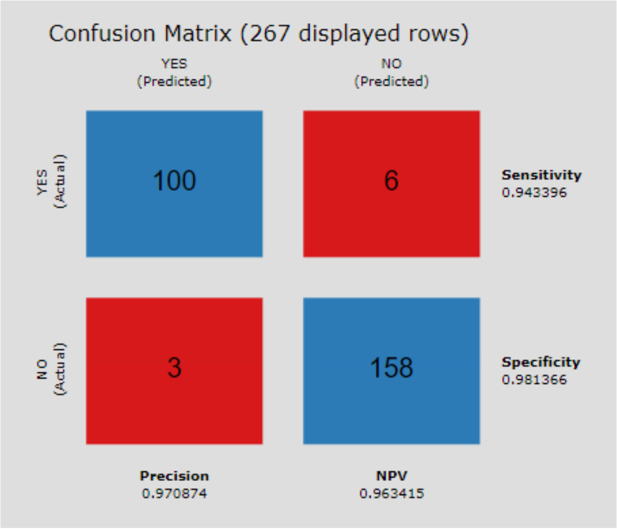


**Supplementary Figure 4S:** Confusion matrix of the test set (N=267) for XGBoost using ErG descriptor. The “YES” label refers to E3 binders, while “NO” refers to non-binder molecules.

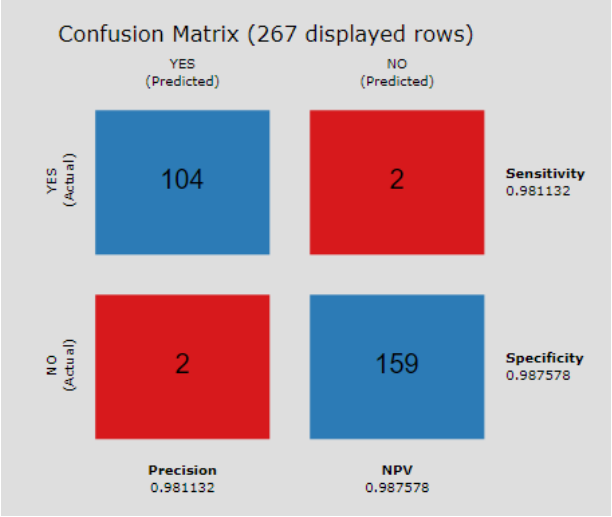


**Supplementary Figure 5S:** Confusion matrix of the test set (N=267) for XGBoost using ChemPhys. The "YES" label refers to E3 binders, while "NO" refers to non-binder molecules

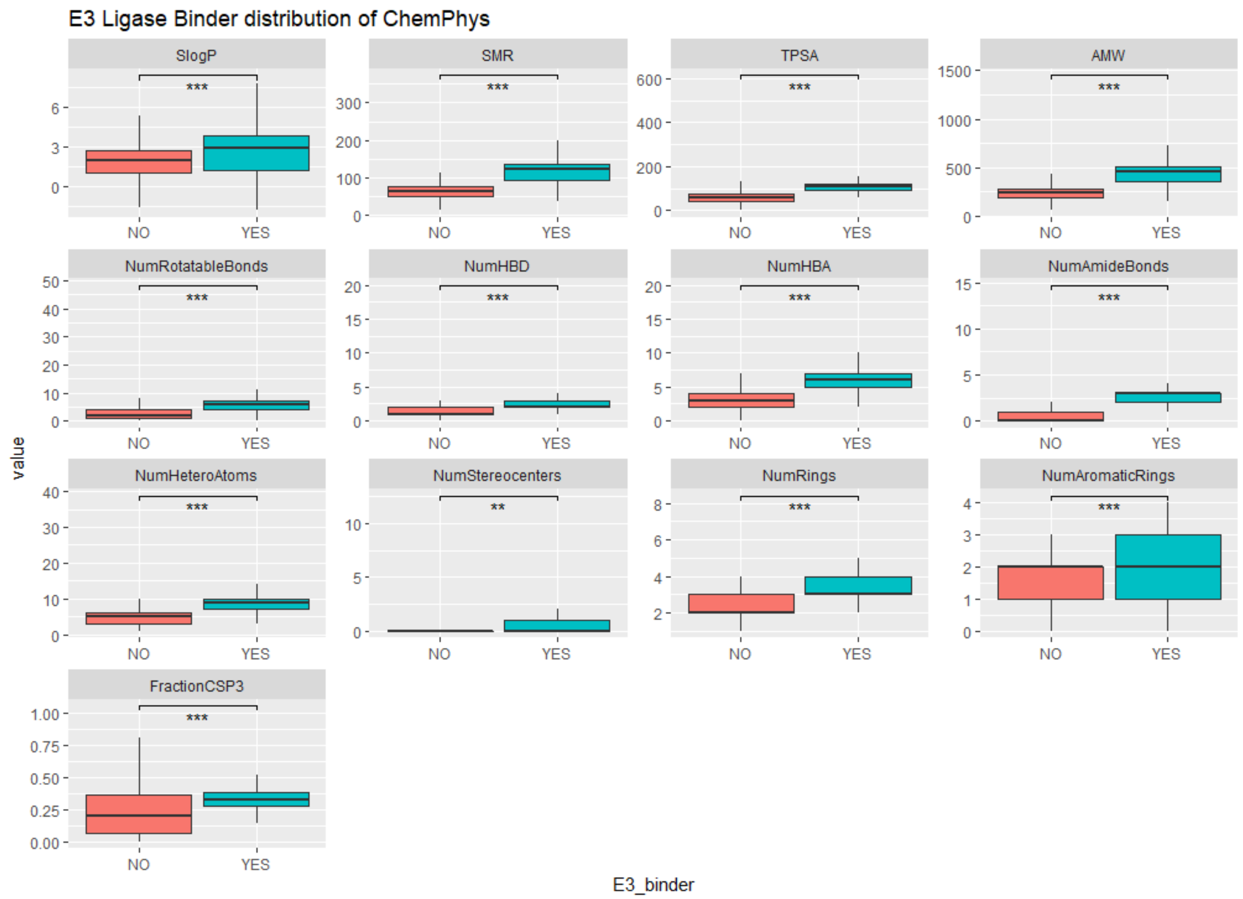


**Supplementary Figure 6S:** Boxplots for ChemPhys description. The 13 chemico-physical descriptors used have been separately analyzed on their distribution through boxplots. Besides the number of stereocenters (** = p-value < 0.1), they all seem to provide significant mean differences (p-value < 0.05 denoted by ***).

| Compound | SMILES | MIA PaCa-2 IC50 (μM) | BxPC-3 IC50 (μM) |
|---|---|---|---|
| Napabucasin | O=C(C)c1oc2C(=O)c3c(C(=O)c2c1)cccc3 | 1.2 ± 0.8 | 1.4 ± 0.3 |
| 1-6 | O=C(NCC[N+]1CCCCC1)c1oc2C(=O)c3c(C(=O)c2c1)cccc3 | 0.1 ± 0.0 | 0.1 ± 0.1 |
| 2-5a | O=C(NCCNc1c2C(=O)N(C(=O)c2ccc1)C1C(=O)NC(=O)CC1)c1oc2C(=O)c3c(C(=O)c2c1)cccc3 | 4.0 ± 0.9 | 1.7 ± 0.7 |
| 2-5b | O=C(NCCCNc1c2C(=O)N(C(=O)c2ccc1)C1C(=O)NC(=O)CC1)c1oc2C(=O)c3c(C(=O)c2c1)cccc3 | 2.7 ± 0.4 | 3.5 ± 0.6 |
| 2-5c | O=C(NCCCCNc1c2C(=O)N(C(=O)c2ccc1)C1C(=O)NC(=O)CC1)c1oc2C(=O)c3c(C(=O)c2c1)cccc3 | 1.60 ± 0.3 | 4.1 ± 1.1 |
| 2-5d | O=C(NCCCCCNc1c2C(=O)N(C(=O)c2ccc1)C1C(=O)NC(=O)CC1)c1oc2C(=O)c3c(C(=O)c2c1)cccc3 | 1.8 ± 0.7 | 3.2 ± 1.3 |
| 2-5e | O=C(NCCCCCCNc1c2C(=O)N(C(=O)c2ccc1)C1C(=O)NC(=O)CC1)c1oc2C(=O)c3c(C(=O)c2c1)cccc3 | 3.0 ± 0.4 | 4.2 ± 1.4 |
| 2-5f | O=C(NCCOCCNc1c2C(=O)N(C(=O)c2ccc1)C1C(=O)NC(=O)CC1)c1oc2C(=O)c3c(C(=O)c2c1)cccc3 | 2.6 ± 0.9 | 4.6 ± 1.2 |
| 2-5h | O=C(NCCOCCOCCOCCNc1c2C(=O)N(C(=O)c2ccc1)C1C(=O)NC(=O)CC1)c1oc2C(=O)c3c(C(=O)c2c1)cccc3 | 2.1 ± 0.8 | 4.5 ± 0.9 |
| 2-5i | O=C(NCCOCCOCCOCCOCCNc1c2C(=O)N(C(=O)c2ccc1)C1C(=O)NC(=O)CC1)c1oc2C(=O)c3c(C(=O)c2c1)cccc3 | 4.4 ± 0.7 | 10.1 ± 3.5 |
| 2-5j | O=C(NCCOCCOCCOCCOCCOCCNc1c2C(=O)N(C(=O)c2ccc1)C1C(=O)NC(=O)CC1)c1oc2C(=O)c3c(C(=O)c2c1)cccc3 | 5.0 ± 1.0 | 7.5 ± 0.4 |
| 3-6a | O=C(NCCN1CC[N+](CCNc2c3C(=O)N(C(=O)c3ccc2)C2C(=O)NC(=O)CC2)CC1)c1oc2C(=O)c3c(C(=O)c2c1)cccc3 | 18.1 ± 11.4 | 25.6 ± 2.0 |
| 3-6b | O=C(NCC[N+]1CCC(CCNc2c3C(=O)N(C(=O)c3ccc2)C2C(=O)NC(=O)CC2)CC1)c1oc2C(=O)c3c(C(=O)c2c1)cccc3 | 1.1 ± 0.3 | 1.1 ± 0.5 |
| 3-6c | O=C(NCC[N+]1CCC(CCCNc2c3C(=O)N(C(=O)c3ccc2)C2C(=O)NC(=O)CC2)CC1)c1oc2C(=O)c3c(C(=O)c2c1)cccc3 | 0.6 ± 0.3 | 0.6 ± 0.1 |
| 4-5 | O=C(NCCCC#Cc1c2C(=O)N(C(=O)c2ccc1)C1C(=O)NC(=O)CC1)c1oc2C(=O)c3c(C(=O)c2c1)cccc3 | 2.1 ± 1.0 | 5.3 ± 0.8 |
| 5-5a | O=C(NCCCCOc1c2C(=O)N(C(=O)c2ccc1)C1C(=O)NC(=O)CC1)c1oc2C(=O)c3c(C(=O)c2c1)cccc3 | 2.0 ± 0.3 | 4.2 ± 0.7 |
| 5-5b | O=C(NCCOCCOCCOc1c2C(=O)N(C(=O)c2ccc1)C1C(=O)NC(=O)CC1)c1oc2C(=O)c3c(C(=O)c2c1)cccc3 | 3.8 ± 1.0 | 8.2 ± 4.1 |
| 6-6a | O=C(NCCOCCOCCNC(=O)c1oc2C(=O)c3c(C(=O)c2c1)cccc3)COc1c2C(=O)N(C(=O)c2ccc1)C1C(=O)NC(=O)CC1 | 16.7 ± 3.8 | 20.4 ± 7.4 |
| 6-6b | O=C(NCCOCCOCCOCCNC(=O)c1oc2C(=O)c3c(C(=O)c2c1)cccc3)COc1c2C(=O)N(C(=O)c2ccc1)C1C(=O)NC(=O)CC1 | 3.7 ± 1.1 | 6.0 ± 5.5 |
| 6-6c | O=C(NCCCCNC(=O)c1oc2C(=O)c3c(C(=O)c2c1)cccc3)COc1c2C(=O)N(C(=O)c2ccc1)C1C(=O)NC(=O)CC1 | 10.5 ± 3.0 | 23.8 ± 3.9 |
| 7-4 | O=C(NCCOCCOCCNc1c2c(C(=O)N(C3C(=O)NC(=O)CC3)C2)ccc1)c1oc2C(=O)c3c(C(=O)c2c1)cccc3 | 13.7 ± 1.0 | 10.5 ± 4.5 |
| 8-1 | O=C(NCCCCCc1c2c(C(=O)N(C3C(=O)NC(=O)CC3)C2)ccc1)c1oc2C(=O)c3c(C(=O)c2c1)cccc3 | 6.4 ± 3.4 | 13.8 ± 7.7 |
| 9-4a | O=C(N[C@@](C(C)(C)C)C(=O)N1[C@](C(=O)NCc2ccc(-c3c(C)ncs3)cc2)C[C@@](O)C1)COCCOCCNC(=O)c1oc2C(=O)c3c(C(=O)c2c1)cccc3 | 14.5 ± 7.1 | >30 |
| 9-4b | O=C(N[C@@](C(C)(C)C)C(=O)N1[C@](C(=O)NCc2ccc(-c3c(C)ncs3)cc2)C[C@@](O)C1)COCCNC(=O)c1oc2C(=O)c3c(C(=O)c2c1)cccc3 | 15.8 ± 4.6 | >30 |
| 12-5 | O=C(NCCOCCOCCNc1cc2C(=O)N(C(=O)c2cc1)C1C(=O)NC(=O)CC1)c1oc2C(=O)c3c(C(=O)c2c1)cccc3 | 3.1 ± 0.4 | 7.9 ± 1.9 |
| 13-11 | O=C(NCC[N+]1CCCCC1)c1oc2C(=O)c3c(C(=O)c2c1)ccc(OCCOCCOCCNc1c2C(=O)N(C(=O)c2ccc1)C1C(=O)NC(=O)CC1)c3 | 3.9 ± 1.1 | 4.2 ± 0.6 |
| XD97 | O=C(NCCOCCOCCNc1c2C(=O)N(C(=O)c2ccc1)C1C(=O)NC(=O)CC1)c1oc2C(=O)c3c(C(=O)c2c1)cccc3 | 1.3 ± 0.5 | 4.3 ± 1.1 |
| XD171 | O=C(NCCOCCOCCNc1c2C(=O)N(C(=O)c2ccc1)C1C(=O)NCCC1)c1oc2C(=O)c3c(C(=O)c2c1)cccc3 | 0.8 ± 0.1 | 2.4 ± 0.9 |
| XD2-149 | O=C(NCC[N+]1CCC(CCCCCNc2c3C(=O)N(C(=O)c3ccc2)C2C(=O)NC(=O)CC2)CC1)c1oc2C(=O)c3c(C(=O)c2c1)cccc3 | 1.0 ± 0.1 | 0.8 ± 0.2 |
| XD2-162 | O=C(NCC[N+]1CCC(CCCCCNc2c3C(=O)N(C(=O)c3ccc2)C2C(=O)N(C)C(=O)CC2)CC1)c1oc2C(=O)c3c(C(=O)c2c1)cccc3 | 2.7 ± 0.4 | 3.4 ± 0.8 |
| Pomalidomide | O=C1C(N2C(=O)c3c(N)cccc3C2=O)CCC(=O)N1 | >30 | >30 |

**Supplementary Table S1:** Table from the supplementary material of paper from Hanafi et al.[1] where all compounds beside Napabucasin were predicted as E3 Ligase binders.
